# The open-closed mod-minimizer algorithm

**DOI:** 10.1101/2024.11.02.621600

**Authors:** Ragnar Groot Koerkamp, Daniel Liu, Giulio Ermanno Pibiri

## Abstract

Sampling algorithms that deterministically select a subset of *k*-mers are an important building block in bioinformatics applications. For example, they are used to index large textual collections, like DNA, and to compare sequences quickly. In such applications, a sampling algorithm is required to select one *k*-mer out of every window of *w* consecutive *k*-mers. The folklore and most used scheme is the *random minimizer* that selects the smallest *k*-mer in the window according to some random order. This scheme is remarkably simple and versatile, and has a *density* (expected fraction of selected *k*-mers) of 2*/*(*w* + 1). In practice, lower density leads to faster methods and smaller indexes, and it turns out that the random minimizer is not the best one can do. Indeed, some schemes are known to approach optimal density 1*/w* when *k → ∞*, like the recently introduced *mod-minimizer* (Groot Koerkamp and Pibiri, WABI 2024).

In this work, we study methods that achieve low density when *k ≤ w*. In this small-*k* regime, a practical method with provably better density than the random minimizer is the *miniception* (Zheng et al., Bioinformatics 2021). This method can be elegantly described as sampling the smallest *closed sycnmer* (Edgar, PeerJ 2021) in the window according to some random order. We show that extending the miniception to prefer sampling *open* syncmers yields much better density. This new method – the *open-closed* minimizer – offers improved density for small *k ≤ w* while being as fast to compute as the random minimizer. Compared to methods based on decycling sets, that achieve very low density in the small-*k* regime, our method has comparable density while being computationally simpler and intuitive.

Furthermore, we extend the mod-minimizer to improve density of any scheme that works well for small *k* to also work well when *k > w* is large. We hence obtain the *open-closed mod-minimizer*, a practical method that improves over the mod-minimizer for all *k*.

## 1 Introduction

Efficient indexing of large textual collections is critical in bioinformatics, and *k*-mer sampling methods play a pivotal role in this [1, 2] as they permit to design sparse, i.e. space-efficient, data structures [3–7]. A popular sampling method is the *minimizer*, simultaneously introduced by Roberts et al. [8] and Schleimer et al. [9]. A minimizer scheme is defined by a triple (*k, w*, 𝒪) and operates as follows: from a window of *w* consecutive *k*-mers of the string to be sampled, the (leftmost) smallest *k*-mer according to the order 𝒪 is elected as the “minimizer” of the window. Since the scheme tends to sample the same *k*-mer from consecutive windows, the set of distinct sampled *k*-mers is a sparse subset of all *k*-mers in the string. Several different sampling algorithms have been proposed in the literature [8–13].

Recently, Groot Koerkamp and Pibiri introduced the *mod-minimizer* [12]. The core idea behind the mod-minimizer is as follows. The position *x* of the smallest *t*-mer in the window is determined and the *k*-mer at position *x* mod *w* is then sampled, where *t* is some small integer parameter (≤ *k*).

This approach is intuitive for several reasons. First, the smallest *t*-mer within the window acts as an “anchor” across potentially many more consecutive windows than the smallest *k*-mer does (hence improving over the random minimizer). If this smallest *t*-mer does not change across these windows, the algorithm exhibits a predictable behavior: either the same *k*-mer is sampled from consecutive windows, or they are spaced apart by exactly *w* positions. This is locally optimal, and indeed it has been mathematically proven that as *k* → ∞ and *w* is fixed, the *density* of the method, that is, the ratio between the expected number of sampled *k*-mers and the total number of *k*-mers in the string, approaches 1*/w*. This effect can be graphically visualized in Figure 1a, that shows an example for (*w, k*) = (4, 31) and *t* = 4. Furthermore, Kille et al. [14] showed that the mod-minimizer has optimal density when *k* ≡ 1 (mod *w*) and the string’s alphabet is large.

**Fig. 1:**
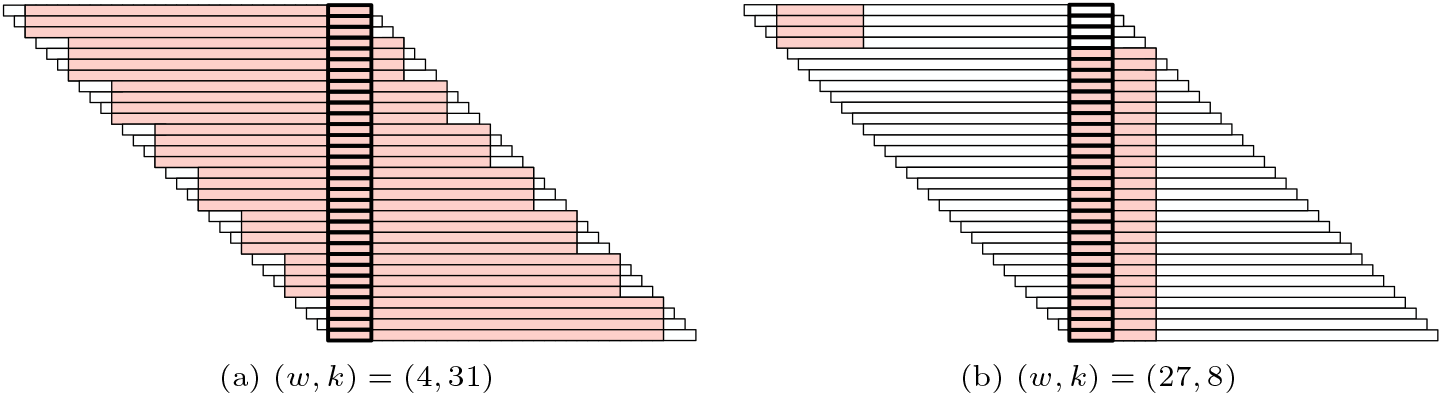
An illustration of the behavior of the mod-minimizer for *w* + *k* − 1 = 34, and *t* = 4. Rows indicate consecutive windows. The thick outlined boxes mark the minimal *t*-mer in each window and the regions highlighted in red indicate the sampled *k*-mer.

**Fig. 2:**
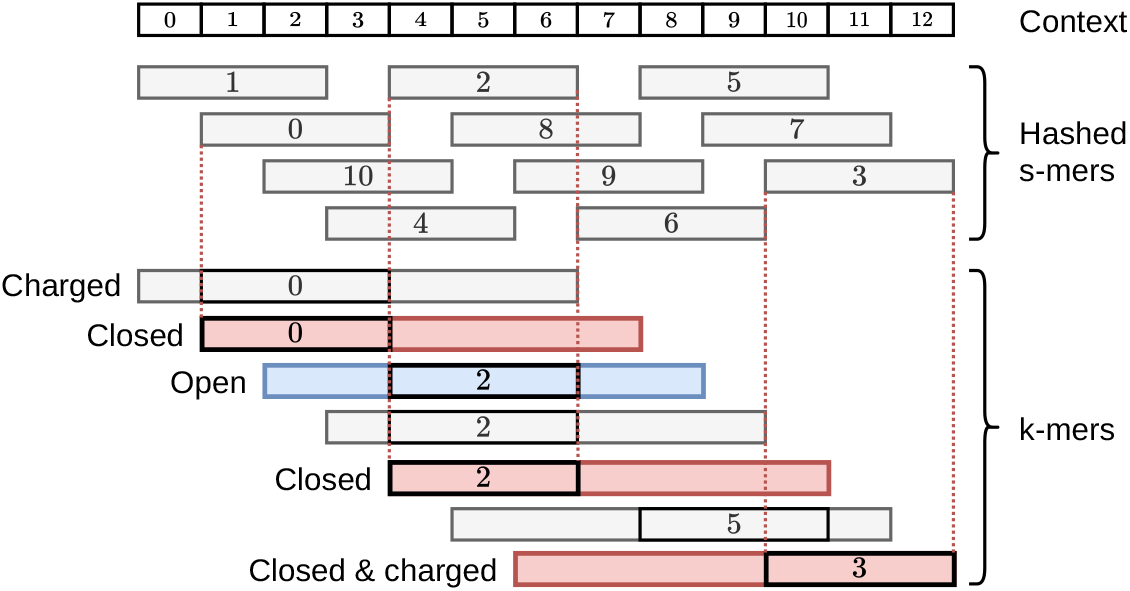
Example of finding all open (blue) and closed (red) syncmers in a context of size *w* + *k*, for *s* = 3, *k* = 7, and *w* = 6. One of the closed syncmers is *charged* because it is the rightmost *k*-mer.

We remark that, regardless of the choice of *t, k* needs to be large compared to *w* for the mod-minimizer to achieve good density. To intuitively see why, let us consider the example in Figure 1b. Let *ℓ* := *w* + *k* − 1 so that an *ℓ*-mer covers *w* consecutive *k*-mers. The example in Figure 1b uses the same value of *ℓ* and the same value of *t* as Figure 1a, hence both figures show a region of an hypothetical string where the smallest *t*-mer is maximally conserved (i.e., for a group of *ℓ* − *t* + 1 consecutive windows) and, as argued above, the behavior of the mod-minimizer is optimal in such region. The two pictures differ only in the choice of the parameters *w* and *k*: a much smaller *k* is used in Figure 1b, namely (*w, k*) = (27, 8). The number of regions of the string where the smallest *t*-mer is maximally conserved, say *M*, only depends on *𝓁* and *t*. Thus, we have 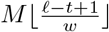 groups of *w* consecutive windows that are optimally sampled and this quantity decreases as *w* increases, like in Figure 1b. This intuitively shows that we cannot use the mod-minimizer with a small value of *k* but we need a different method.

### Contributions

This work introduces the *open-closed mod-minimizer* – a sampling algorithm that has lower density than the best known schemes for *k > w*, like the mod-minimizer, and also generally works for *any* value of *k* (Figures 3 and 5). This new scheme is achieved by combining two main ingredients:

**Fig. 3:**
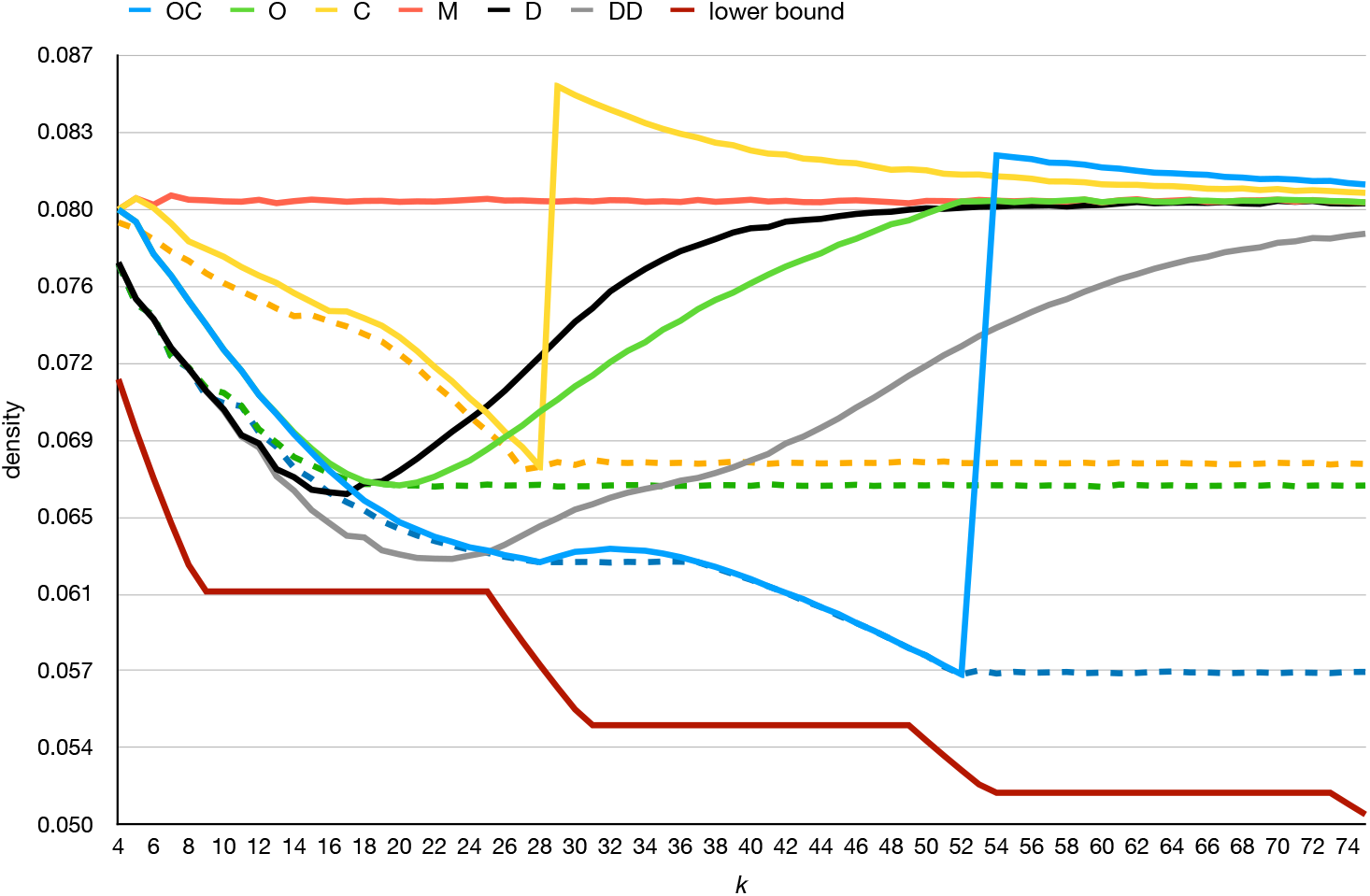
Density for *w* = 24 and varying *k*, measured on a random string of ten million i.i.d. random characters for *σ* = 4. For the methods `OC, O`, and `C`, we use *s* = 4 for the solid lines. The dashed lines, instead, use the best choice of *s*.

**Fig. 4:**
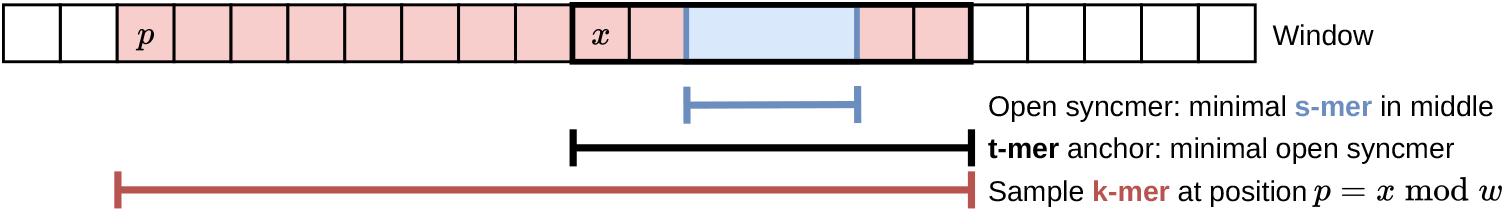
Example of the open-closed mod-minimizer for *s* = 3, *t* = 7, *k* = 15, and *w* = 8.

**Fig. 5:**
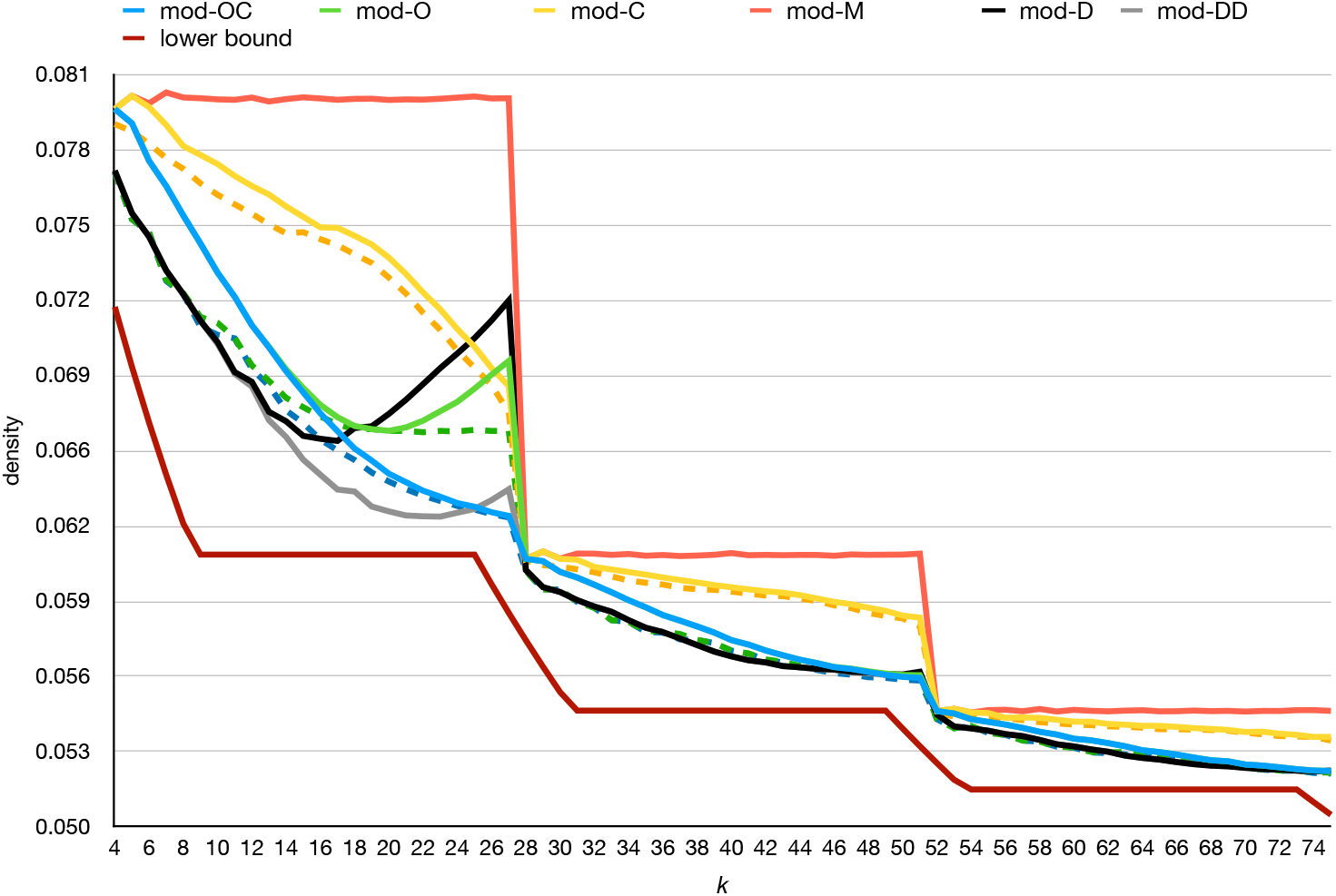
Density for *w* = 24 and by varying *k*, measured on a random string of ten million i.i.d. random characters for *σ* = 4. We use *r* = 4 for all methods. For the methods mod-OC, mod-O, and mod-C, we use *s* = 4 for the solid lines. The dashed lines, instead, use the best choice of *s*.

#### 1. The open-closed minimizer

Among the methods that improve the random minimizer for *k* ≤ *w*, the *miniception* by Zheng et al. [10] stands out for its elegance: it samples the smallest *closed syncmer* according to some random order. (A closed syncmer is a *k*-mer whose smallest substring of length *s* ≤ *k* appears either at the beginning or at the end [15].) We extend the miniception to also consider *open* syncmers – *k*-mers whose smallest contained *s*-mer is in the middle position [15, 16]. Specifically, the smallest open syncmer in the window is preferred, with respect to a random order on *k*-mers. If there are none, the smallest closed syncmer is preferred. Lastly, if no closed syncmer is present either, the smallest *k*-mer is considered. We show that this method – that we name the *open-closed minimizer* – significantly improves the density of the miniception to be comparable with the *double decycling* method of Pellow et al. [11], making it a practical and useful method when *k* ≤ *w*.

As an example application, we used the open-closed minimizer as a replacement of the random minimizer in SSHash [3, 4], a recent *k*-mer dictionary based on minimizers. For default parameters (*w, k*) = (11, 21), the open-closed minimizer consistently yields SSHash indexes that are 14% smaller across several datasets.

#### 2. The extended mod-minimizer

We then generalize the mod-minimizer [12] to accommodate any anchoring mechanism, not just the random minimizer on *t*-mers. In this way, the mod-minimizer can be seen as a general method to improve the density of any scheme for *k > w*. In particular, instead of using the *smallest t*-mer, one could use the *open-closed minimizer* of length *t* as introduced above, hence obtaining the so-called *open-closed mod-minimizer*. Furthermore, the new schemes obtained by this extended mod-minimizer framework retain the benefit of being computationally efficient, making them practical for large-scale applications.

Again, for parameters (*w, k*) = (11, 21), the open-closed mod-minimizer makes SSHash 18% smaller when indexing the whole human genome (GRCh38), reducing space usage from 8.70 bits/*k*-mer to 7.13 bits/*k*-mer.

One drawback is that a formal analysis of the density of the extended mod-minimizer tends to be more difficult than for the mod-minimizer based on random minimizers.

### Software

Both C++ and Rust implementations of the proposed algorithms are publicly available on GitHub at

- https://github.com/jermp/minimizers, and
- https://github.com/RagnarGrootKoerkamp/minimizers, respectively

### Organization

The rest of the article is organized as follows. Section 2 fixes the notation used throughout the article and gives preliminary definitions. In Section 3, we study the case where *k* ≤ *w* and introduce an open-closed minimizer. In Section 4, we extend the mod-minimizer to improve the density of the methods discussed in Section 3 to *k > w*, culminating in the open-closed mod-minimizer. We conclude in Section 5 where we also discuss some promising future work.

Experimental results are presented directly in Section 3 and Section 4 respectively, instead of postponing them to the end of the paper. We believe this is a good way to guide the reader through solutions of incremental sophistication. Thus, we report here some details about our experimental setup.

For all experiments we use the C++ implementation of the algorithms, compiled with gcc 11.1.0 under Ubuntu 18.04.6. Whenever we need to hash *k*-mers, we use the 128-bit pseudorandom hash function CityHash [17]. We do not explicitly consider the time required to compute the methods (i.e., to sample the *k*-mers from a string) because all of them can be implemented efficiently, and thus have comparable runtime.

## 2 Preliminaries

Following Groot Koerkamp and Pibiri [12], we here fix some preliminary notions and precisely define the problem under study. Table 1 summarizes the most common notation that we will use throughout the paper.

**Table 1:**
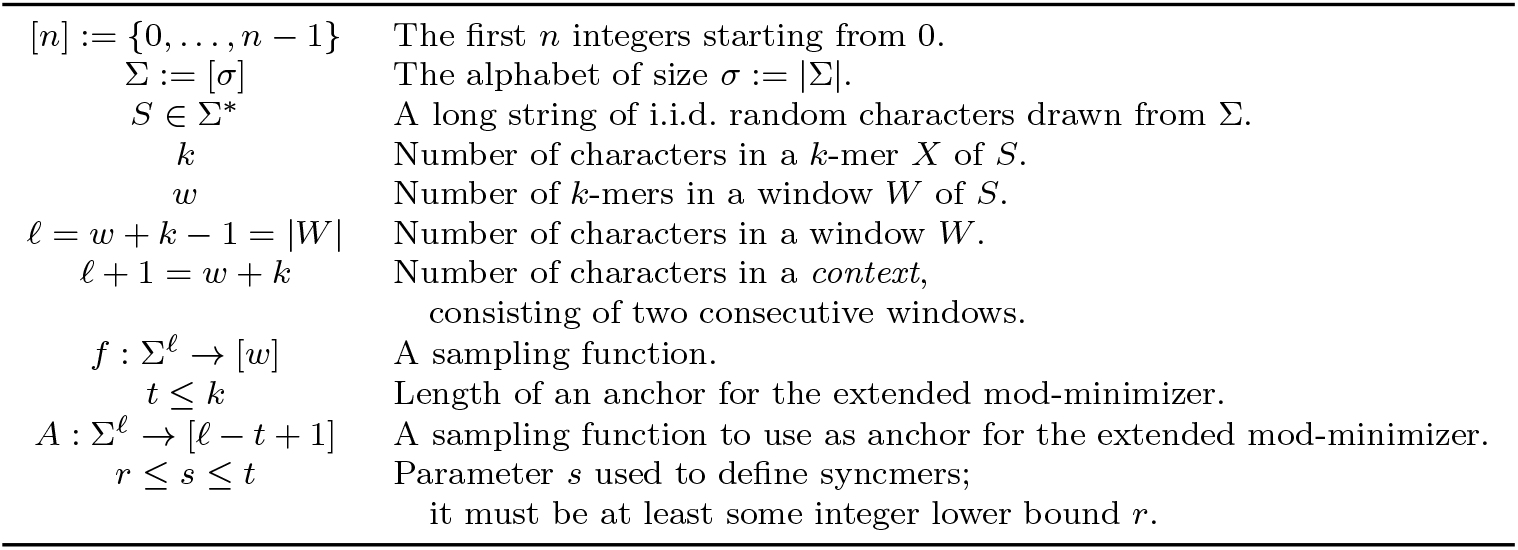
Notation used in this article.

**Table 2:** Space usage for SSHash indexes in bits/*k*-mer across different datasets and for the minimizer types proposed in this article. The used parameters are (*w, k*) = (11, 21) for all datasets. We show percentages relative to the random minimizer, which is the default option to build SSHash indexes.

| Minimizer | Human Chr. 13 | Whole human | Salmonella-100 | Axolotl |
| --- | --- | --- | --- | --- |
| random | 7.53 | 8.70 | 7.55 | 9.91 |
| open-closed (Section 3.1) | 6.45<br>(−14.35%) | 7.44<br>(−14.48%) | 6.69<br>(−11.39%) | 8.54<br>(−13.80%) |
| open-closed mod (Section 4.1) | 6.18<br>(−17.90%) | 7.13<br>(−18.05%) | 6.46<br>(−14.42%) | 8.22<br>(−17.05%) |

### Basic notation

Let [*n*] := {0, …, *n* − 1}, for any *n* ∈ ℕ. We fix an alphabet Σ = [*σ*] of size *σ* = 2^*O*(1)^. Let *S* ∈ Σ^*^ be a string. We refer to *S*[*i*..*j*) as its sub-string of length *j* − *i* starting at index *i* and ending at index *j* (excluded). When *j* − *i* = *k* for some *k* ≥ 1, we call *S*[*i*..*j*) a *k*-mer of *S*. In the following, let *w >* 0 be an integer, so that any string of length *ℓ* = *w* + *k* − 1 defines a *window W* of *w* consecutive *k*-mers. Each *k*-mer in *W* can be uniquely identified with an integer in [*w*], corresponding to its starting position in *W*. A window is always implicitly assumed to be a substring of a hypothetical long string *S*. We say that two windows *W* and *W*^*′*^ are *consecutive* when *W* [1..*ℓ*) = *W*^*′*^[0..*ℓ* − 1).

We write *a* mod *m* for the remainder of *a* after division by *m* and *a* ≡ *b* (mod *m*) to say that *a* and *b* have the same remainder modulo *m*.

### Orders and hashes

An order 𝒪_*k*_ on *k*-mers is a function 𝒪_*k*_ : ∑^*k*^ → ℝ, such that 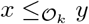 if and only if 𝒪_*k*_(*x*) ≤ 𝒪_*k*_(*y*). We do not necessarily require 𝒪_*k*_ to be *random*, although practitioners often use a (pseudo-)random hash function *h* : ∑^*k*^ → [*U*] to define the order, where [*U*] is a sufficiently large range, like *U* = 2^128^. We therefore make the standard assumption [18, 19] that *h* is drawn from a family of fully random hash functions that can be evaluated in *O*(1) on a machine word. Unless otherwise specified, all the orders we consider in this work are random.

Since *σ* = 2^*O*(1)^, it follows that any *k*-mer *x* ∈ ∑^*k*^ fits in *O*(*k*) words and *h*(*x*) is computed in *O*(*k*) time. Furthermore, using a rolling hash function [20], we can compute *w* hashes for the *w* consecutive *k*-mers in a window in *O*(*w* + *k* − 1) rather than the naive *O*(*wk*). We implicitly assume this linear bound when discussing the complexities of the algorithms.

### Sampling functions and their densities

All methods we consider in this article can be expressed as a function *f* : ∑^*w*+*k−*1^ → [*w*] that, given a window *W*, samples the *k*-mer starting at position *f* (*W*) in *W*. We call such function *f* a *sampling* function and, sometimes, we colloquially refer to *f* as a “scheme”.

What is a “good” scheme? The performance metric we focus on in this work is the *density* of a scheme, defined as follows. Given a string *S* of length *n*, let *W*_*i*_ := *S*[*i*..*i*+*ℓ*) for *i* ∈ [*n* − *ℓ* + 1]. A sampling function *f* selects the *k*-mers starting at positions *i* + *f* (*W*_*i*_) *i* [*n ℓ* + 1]. The *particular density* of *f* on *S* is |{ *i* + *f* (*W*_*i*_) | *i* ∈ [*n* − *ℓ* + 1] */* (*n* − *k* + 1). The *density* of *f* is defined as the expected particular density on a string *S* consisting of i.i.d. random characters of ∑ in the limit where *n* → ∞. We remark that, in practice, we use a finite but sufficiently-long random string *S* to approximate the density of a scheme in this work.

### Problem statement

With these initial remarks in mind, we can state precisely the problem addressed in this article.

#### Problem 1

(Pure sampling function problem). *Given integers w* ≥ 2 *and k* ≥ 1, *implement a function f* : ∑^*w*+*k−*1^ → [*w*] *in O*(1) *space with as low density as possible*.

## 3 The small-*k* case: the open-closed minimizer

In this section, we study methods that perform well when *k* ≤ *w*. We refer to this case as the “small-*k*” case. This scenario is particularly relevant to implement, e.g., sparse data structures for *ℓ*-mers [3–5] and building of De Bruijn graphs [21–23] just to mention two example applications. In these applications, the high-level idea is to

### Algorithm 1

Pseudocode for the random minimizer algorithm.

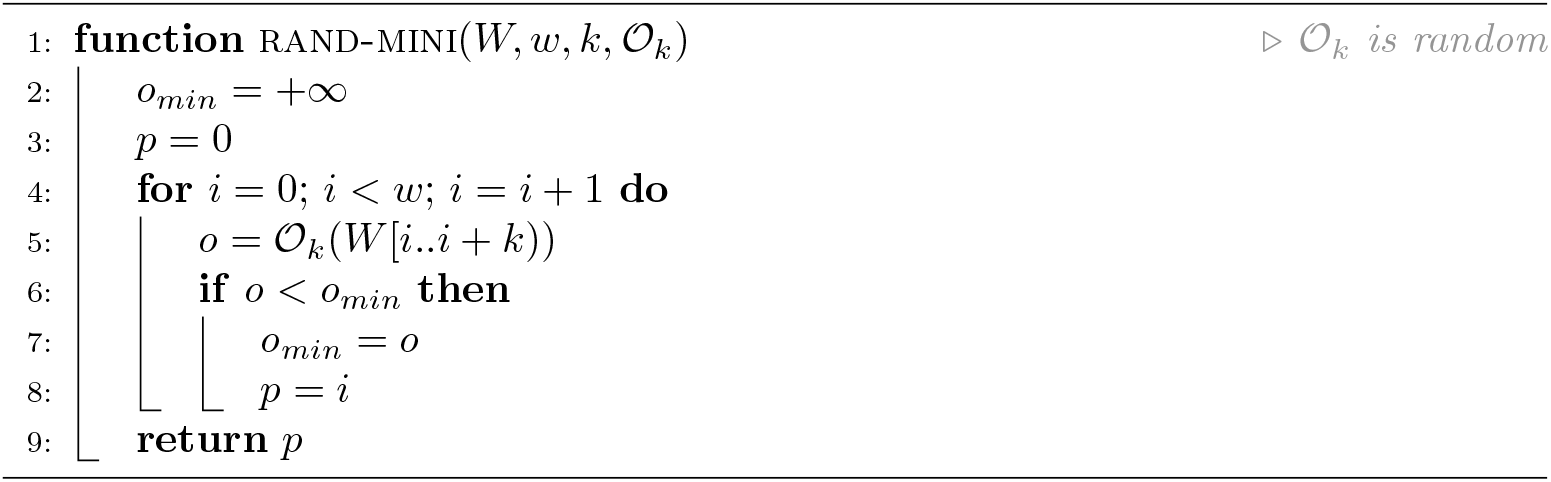

“cluster” similar *ℓ*-mers together to accelerate queries and improve compression. In particular, this is done using minimizers, as follows. For each *ℓ*-mer, its minimizer is computed, and all *ℓ*-mers having the same minimizer belong to the same cluster. In these cases, *k* is typically fixed (e.g., *k* = 20) and *w* increases, so that usually *k* is less than *w*.

### 3.1 The open-closed minimizer

The “classic” random minimizer is the simplest minimizer algorithm: it selects the smallest *k*-mer of the window according to some random order 𝒪_*k*_. (We remind the reader that all orders used in this article are random.) If two or more *k*-mers have the same smallest rank, then the leftmost *k*-mer is considered. Computing a minimizer thus takes *O*(*w* + *k* − 1) time and it has density 2*/*(*w* + 1) + *o*(1*/w*) (when *k* is not too small^1^) [10]. Efforts spent in improving its density spurred many research results [10, 12, 13, 25–28] and it is the default choice in practical algorithm engineering [3–7, 21, 23]. For these reasons, we give the corresponding pseudocode in Algorithm 1.

Among the methods that perform better than a random minimizer, the *miniception* by Zheng et al. [10] has provably lower density, even when *k* ∼ *w*. The miniception can be elegantly described in terms of *closed syncmers*. Hence we first describe those.

#### Closed and open syncmers

The definitions of closed and open syncmers were first given by Edgar [15]. For a given parameter 1 ≤ *s* ≤ *k*, a *k*-mer is a *closed syncmer* if its smallest contained *s*-mer is either in first or last position, i.e., in position 0 or *k* − *s* respectively. Note that this definition is *context free*, in that whether a *k*-mer is a closed syncmer does not depend on surrounding characters. Closed syncmers satisfy a window guarantee of *k* − *s*, meaning that there is at least one closed syncmer in any window of *w* ≥ *k* − *s* consecutive *k*-mers. Closed syncmers have a density of 2*/*(*k* − *s* + 1) (assuming a random order on *s*-mers), which is the same as that of a random minimizer when *s* = *k* − *w* for *k > w*. Indeed, syncmers were designed to improve the *conservation* metric rather than density compared to minimizers (see the original paper by Edgar [15] for details).

A variation on the closed syncmer is the *open syncmer*, where the smallest contained *s*-mer is required to be at a specified offset *v* ∈ [*k* − *s* + 1]. Shaw and Yu [16] showed that choosing *v* = ⌊ (*k* − *s*)*/*2⌋ is best for conservation, so we assume this choice too. Unlike closed syncmers, open syncmers have a *distance guarantee*: two consecutive open syncmers are always at least ⌊(*k* − *s*)*/*2⌋ + 1 positions apart.

##### Algorithm 2

Pseudocode for the *miniception* (left) and the *open-closed* (“OC”, right) minimizer methods. The differences are highlighted in blue.

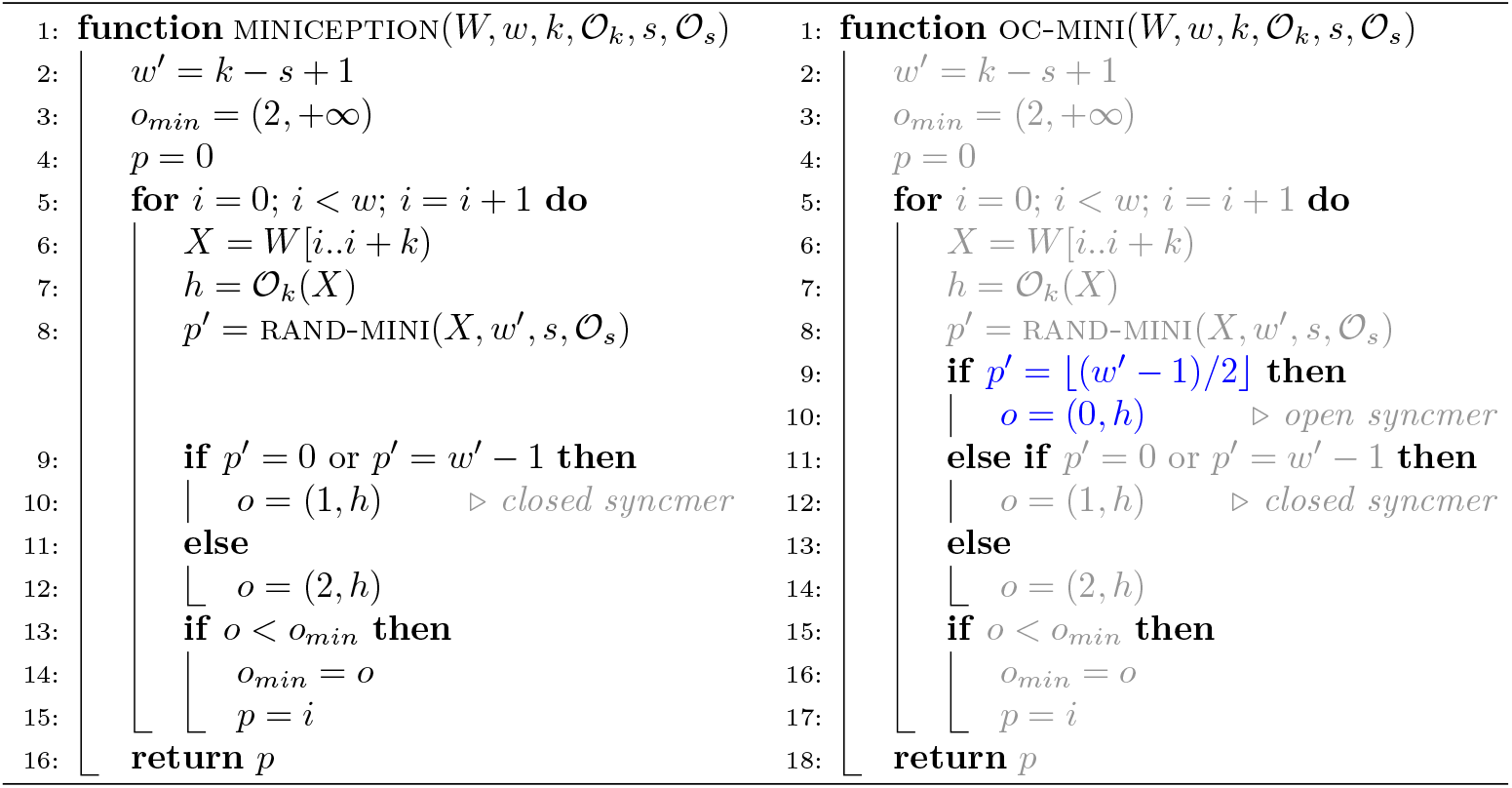

#### The miniception

With these definitions in mind, it is easy to describe the miniception. The term *miniception* stands for “minimizer inception” and the method samples the smallest closed syncmer from the window, according to a random order 𝒪_*k*_. Algorithm 2 (left) illustrates the method.

At a high level, the idea of this method (as well as other methods, e.g., the decycling set based method by Pellow et al. [11]) is to first use a context-free scheme to sample some fraction of *k*-mers. Then, in windows with none or multiple sampled *k*-mers, a random order is used as a tiebreaker. It is clear that such schemes fall back to a random minimizer when almost all or almost no *k*-mers are sampled. Thus, intuitively, they perform best when the sampled fraction is on the order of 1*/w*.

Note that when *w < k* − *s*, a window may not contain any closed syncmer. In fact, while positions {0, …, *w* − 1} and {*k* − *s*, …, *k* − *s* + *w* − 1} of the smallest *s*-mer in a window induce closed syncmers respectively at positions {0, …, *w* − 1}, the *k* − *s* − *w* positions in between these two sets, i.e., {*w*, …, *k* − *s* − 1}, do not induce any closed syncmers. Note that a *s*-mer starting at any of such “middle” positions is a substring of all *k*-mers in the window. Thus, if it is the smallest in the window, it is also the smallest for all *k*-mers, preventing all *k*-mers to be closed syncmers. Assuming no duplicate *s*-mers in a window and a random order _*s*_, the probability that a window contains no closed syncmers is therefore 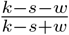 for *w < k* − *s*. This is the probability with which the miniception samples the smallest *k*-mer.

#### The open-closed minimizer

Inspired by the miniception, we here propose a natural extension of the method to *open* syncmers. The method is illustrated in Algorithm 2 (right), with the differences to the miniception highlighted. Specifically, the open-closed minimizer prefers sampling the smallest open syncmer (line 9-10). If no open syncmer is found in the window, then the smallest closed syncmer is considered (line 11-12), like in miniception. Lastly, if no closed syncmer is found either, the smallest *k*-mer is sampled. We call this new method of sampling syncmers the “*open-closed minimizer* “.

The rationale behind this method is that open syncmers have a distance *lower* bound, i.e., we know that two consecutive open syncmers must be at least ⌊(*k* − *s*)*/*2⌋ +1 positions apart. This is in contrast to closed syncmers that do not have a similar guarantee (but instead have an *upper* bound on the distance between them). As it turns out, the distance lower bound of open syncmers should be preferred over closed syncmers.

As already discussed for the miniception, the smallest *s*-mer positions {0, …, *w* − 1} and {*k* − *s*, …, *k* − *s* + *w* − 1} induce closed syncmers. Further, the positions *i* in the middle, i.e., *i* ∈ {*w*, …, *k* − *s* − 1}, induce an open syncmer when 0 *i* ⌊(*k s*)*/*2⌋ *< w*. From this, we can infer that it is possible that no open nor closed syncmer is present in a window when *w <* (*k* − *s*)*/*2 or equivalently, *k >* 2*w* + *s*.

### 3.2 Analysis

To obtain the exact density of the open-closed minimizer, like for any other forward scheme, one could compute the number of sampled *k*-mers on a De Bruijn sequence of order *w* + *k* (a cyclic string where each possible sub-string of length *w* + *k* occurs once) [29]. However, this takes exponential time as the sequence has length *σ*^*w*+*k*^, and thus quickly becomes infeasible. Here, we present a polynomial method to compute the density.

Instead, it is possible to consider a *context* of *w* + *k* characters, containing two consecutive windows. The density then equals the probability that the two windows sample a different *k*-mer. For minimizer schemes specifically, this corresponds to the probability that either the first *k*-mer (that in position 0) or the last *k*-mer in the context (that in position *w*) is sampled [9, 10]. For convenience, we will call these two *k*-mers at the edges *charged k-mers*.

Let us consider some simple examples before presenting the case of the open-closed minimizer. We assume that all *k*-mers and *s*-mers in a window are distinct.

**Example 1: the random minimizer**. For the random minimizer, there are always *w* + 1 *k*-mers in the context of length *w* + *k* among which to pick the smallest one, hence the probability that the context is charged is 2*/*(*w* + 1), assuming all *k*-mers are distinct.

**Example 2: the miniception**. Now, generalizing it to the miniception, first we have to count the number of closed syncmers in a context. Call this quantity *C*. Naturally we have 0 ≤ *C* ≤ *w* + 1. Note that when *C* = 0, i.e., there are no closed syncmers in the context, then miniception “falls back” to random minimizers and thus the context is charged with probability 2*/*(*w* + 1). Assume now that *C >* 0. Among those *C k*-mers that are closed syncmers, let *C*_*c*_ be the number of charged ones, i.e., those at position 0 or *w* in the context. Clearly, 0 ≤ *C*_*c*_ ≤ min(2, *C*). The probability that the context is charged is then *C*_*c*_*/C*. Of course, not all configurations (*C, C*_*c*_) are equally probable. For example, it is far more likely to have 2 closed syncmers in a context rather than *w* + 1. Therefore, we would like to compute the probability distribution of the count configurations (*C, C*_*c*_), i.e., ℙ{a context has configuration (*C, C*_*c*_)} for all (*C, C*_*c*_). The density of miniception is then

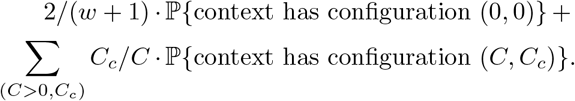

Table 3 in Appendix A shows the distribution of the configurations (*C, C*_*c*_) for *w* = 5, *k* = 11, and *s* = 6, under the assumption that there are no duplicate *s*-mers in a context. In this case, the computed density is 0.2929, which exactly matches what is measured in practice over a long random string.

**Table 3:** Probability distribution of the configurations (*C, C*_*c*_) (as used, e.g., by miniception [10]) for *w* = 5, *k* = 11, and *s* = 6. In this case, the computed density is 0.2929.

| $(C, C_c)$ | $\mathbb{P}\{\text{context has configuration } (C, C_c)\}$ | $\mathbb{P}\{\text{context is charged}\}$ |
| --- | --- | --- |
| (0, 0) | 0.0 | 0.33 |
| (1, 0) | $2.65 \times 10^{-1}$ | 0.0 |
| (1, 1) | 0.0 | 1.0 |
| (2, 0) | $1.76 \times 10^{-1}$ | 0.0 |
| (2, 1) | $2.31 \times 10^{-1}$ | 0.5 |
| (2, 2) | $1.11 \times 10^{-1}$ | 1.0 |
| (3, 0) | $4.11 \times 10^{-2}$ | 0.0 |
| (3, 1) | $1.09 \times 10^{-1}$ | 0.33 |
| (3, 2) | $2.56 \times 10^{-2}$ | 0.67 |
| (4, 0) | $3.37 \times 10^{-3}$ | 0.0 |
| (4, 1) | $2.25 \times 10^{-2}$ | 0.25 |
| (4, 2) | $1.21 \times 10^{-2}$ | 0.5 |
| (5, 0) | 0.0 | 0.0 |
| (5, 1) | $1.73 \times 10^{-3}$ | 0.2 |
| (5, 2) | $2.50 \times 10^{-3}$ | 0.4 |
| (6, 0) | 0.0 | 0.0 |
| (6, 1) | 0.0 | 0.17 |
| (6, 2) | $1.92 \times 10^{-4}$ | 0.33 |

#### The open-closed minimizer

Now, to extend the analysis to the open-closed minimizer, we have to also take into account the number of open syncmers, say *O*, and the number of those that are charged in a context, say *O*_*c*_. In other words, we have to compute the probability distribution of the count configurations (*O, C, O*_*c*_, *C*_*c*_). Note that as soon as there is at least one open syncmer (*O >* 0), then the counts (*C, C*_*c*_) are irrelevant for computing the density. Table 4 in Appendix A shows the distribution of the configurations (*O, C, O*_*c*_, *C*_*c*_) for *w* = 5, *k* = 11, and *s* = 6, where we omit for conciseness the configurations whose probability is 0. In this case, the computed density is 0.2864.

**Table 4:**
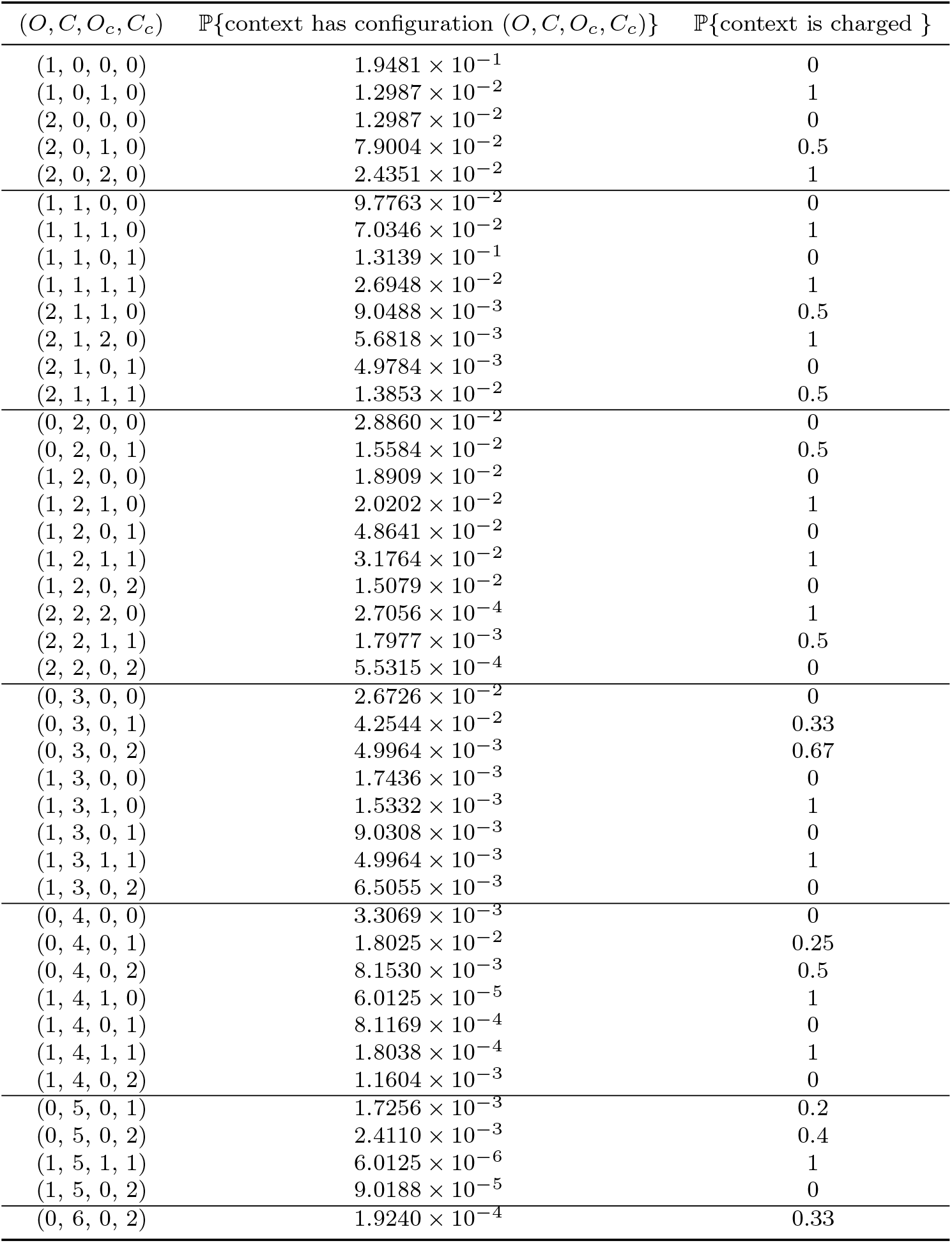
Probability distribution of the configurations (*O, C, O*_*c*_, *C*_*c*_) (as used by the open-closed minimizer from Section 3.1) for *w* = 5, *k* = 11, and *s* = 6. In this case, the computed density is 0.2864.

#### Computing the probability distribution: brute force

We now address the problem of computing the probability distribution of the configurations (*O, C, O*_*c*_, *C*_*c*_). One straightforward way to do so is to consider each possible permutation of (the hashes of) the *s*-mers in a context and derive the corresponding configuration (*O, C, O*_*c*_, *C*_*c*_).

Let us consider an example. Assume *w* = 6, *k* = 7, and *s* = 3, so that there are *w* + *k* − *s* + 1 = 11 distinct *s*-mers in a context. To infer an order between the *s*-mers we can think of each of them as having a distinct hash. For example, assume that the hashes are [1, 0, 10, 4, 2, 8, 9, 6, 5, 7, 3], as in Figure 2. This order induces three closed syncmers, at (zero-based) positions 1, 4, and 6. Specifically, we have a closed syncmer at position 1 because among the *k* − *s* + 1 = 5 *s*-mers within the *k*-mer at position 2 in the context, i.e., with hashes [0, 10, 4, 2, 8], the smallest *s*-mer (underlined) is in the first position. Among those 3 closed syncmers, the one at position 6 is charged. There is a single (uncharged) open syncmer at position 2, since it contains *s*-mers with hashes [10, 4, 2, 8, 9], the smallest of which is in the middle, at offset ⌊(*k* − *s*)*/*2⌋ = 2. Summing up, the configuration of the context is (*O, C, O*_*c*_, *C*_*c*_) = (1, 3, 0, 1).

By enumerating and analyzing all possible orders of *s*-mers in this way, we keep track of how many orders have configuration (*O, C, O*_*c*_, *C*_*c*_), say *N*, and compute ℙ {context has configuration (*O, C, O*_*c*_, *C*_*c*_) } as *N/*(*w* + *k* − *s* + 1)!. As the number of orders to consider is (*w* +*k − s*+1)!, this approach is feasible for only very small values of *w* and *k*.

#### Computing the probability distribution: recursion

We now introduce a recursive method to compute the probability distribution of the configurations (*O, C, O*_*c*_, *C*_*c*_) and, hence, the density of open-closed minimizers, that runs in time polynomial in the number of *s*-mers in a context, *w* + *k − s* + 1. We assume that there are no duplicate *s*-mers in a context. Pseudocode is shown in Algorithm 3.

The method first considers the position of the smallest *s*-mer. Since the order on *s*-mers is random, this position *i* is uniform in {0, …, *w* + *k* − *s*}. Once the smallest *s*-mer is known, we can determine for all *k*-mers containing the *s*-mer whether they are a (charged) open or closed syncmer. Further, the smallest *s*-mer splits the remaining *k*-mers into those on the left and right of it. These two groups are independent of each other: the probability that a *k*-mer left of the *s*-mer is a syncmer is independent of a *k*-mer right of the *s*-mer being a syncmer. This allows us to recurse on these two halves independent from each other. We then add (the probability distributions of) the counts of the left and right part, and take the average over all choices of *i*.

Now, consider the recursion in more detail. First, a range of less than *k* characters can not contain any syncmers, and hence has probability 1 for counts (0, 0, 0, 0). Otherwise, consider the position 0 ≤ *i* ≤ *w* + *k* − *s* of the smallest *s*-mer. Then, one of the following three events can happen:

1. When *i* is sufficiently far away from the boundaries, the *k*-mer containing the minimal *s*-mer as its middle *s*-mer is fully contained in the range, and hence is an open syncmer. When this *k*-mer is the first or last in the window, we additionally count it as a charged open syncmer.
2. Otherwise, we consider closed syncmers. If we can extend the chosen *s*-mer left and/or right by *k* − *s* characters, then those (up to two) *k*-mers are closed syncmers. And as before, we also count how many of the two are charged.

#### Algorithm 3

Pseudocode to compute the density of the open-closed (mod-) minimizer in polynomial time. For the open-closed minimizer, simply set *t* = *k*. To compute the density of the closed minimizer (a.k.a., miniception) or open minimizer, ignore the if statements at line 6 or 8.

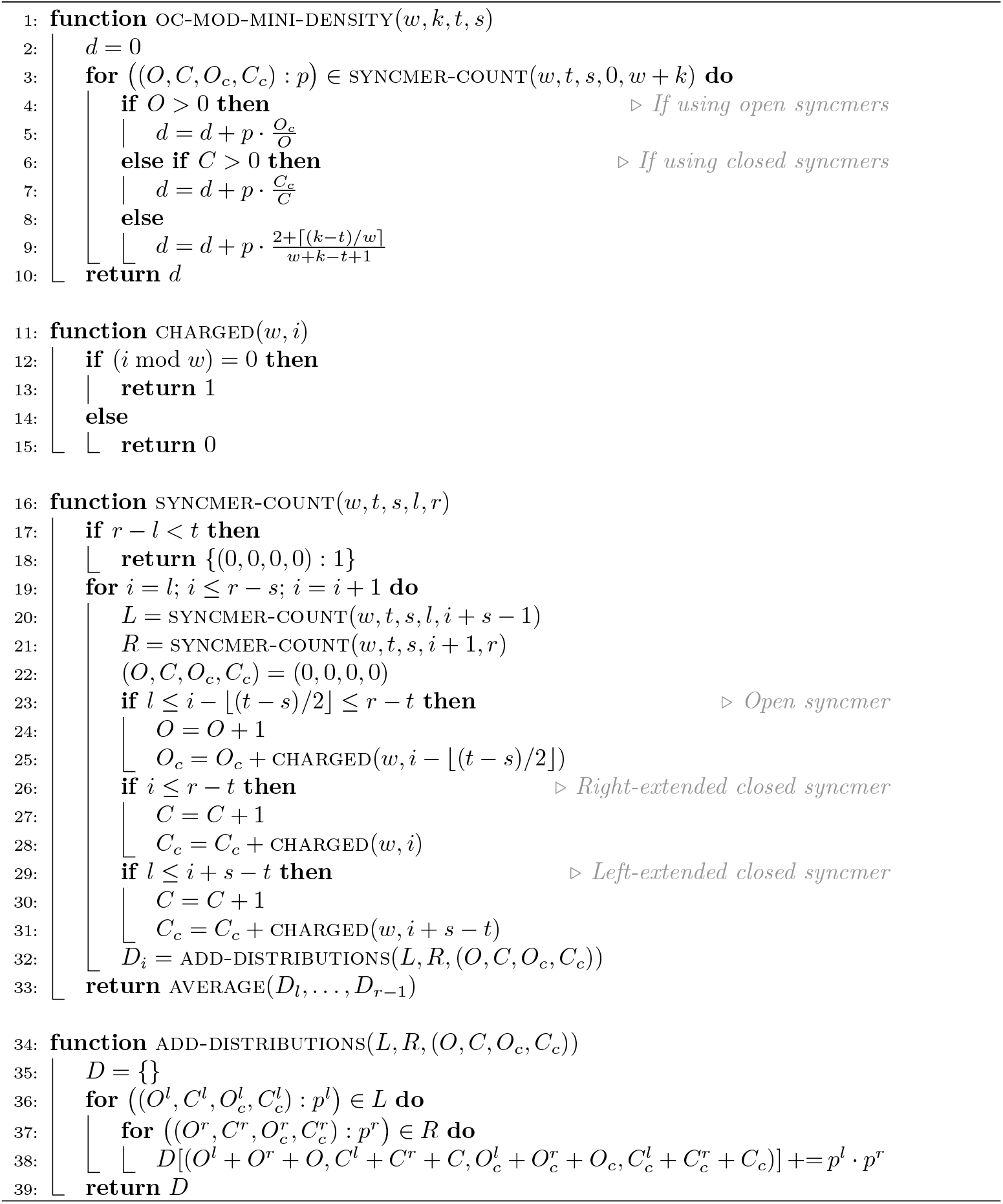
3. If the chosen *s*-mer does not induce an open or closed syncmer, we simply do not increase the counts.

After counting the open/closed syncmers containing the *s*-mer at position *i*, we use recursion to count the number of open/closed syncmers in *W* [0..*i* + *s* − 1) and *W* [*i* + 1..*w* + *k*), with the modification that for the recursive steps, the leftmost and/or rightmost *k*-mer in the remaining interval may not be the leftmost/rightmost *k*-mer in the full window, and hence not be charged.

In conclusion, the algorithm described here can be used to compute exactly the density of the open-closed minimizer. However, we lack a closed-form formula for its density or a tight approximation. We leave these two problems for the future.

### 3.3 Density

Figure 3 compares the density of the described schemes for *w* = 24 and by varying *k*, over a string of ten million i.i.d. random characters drawn from alphabet of size *σ* = 4 (we choose this value of *σ* as it is used when sampling DNA sequences; Figure 6a in Appendix B shows the same plot for *σ* = 256). The curve named “lower bound” corresponds to the (simplified) lower bound proved by Kille et al. [14], which is 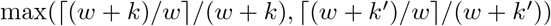 where *k*^*′*^ is the smallest integer ≥ *k* such that *k*^*′*^≡ 1 (mod *w*).

**Fig. 6:**
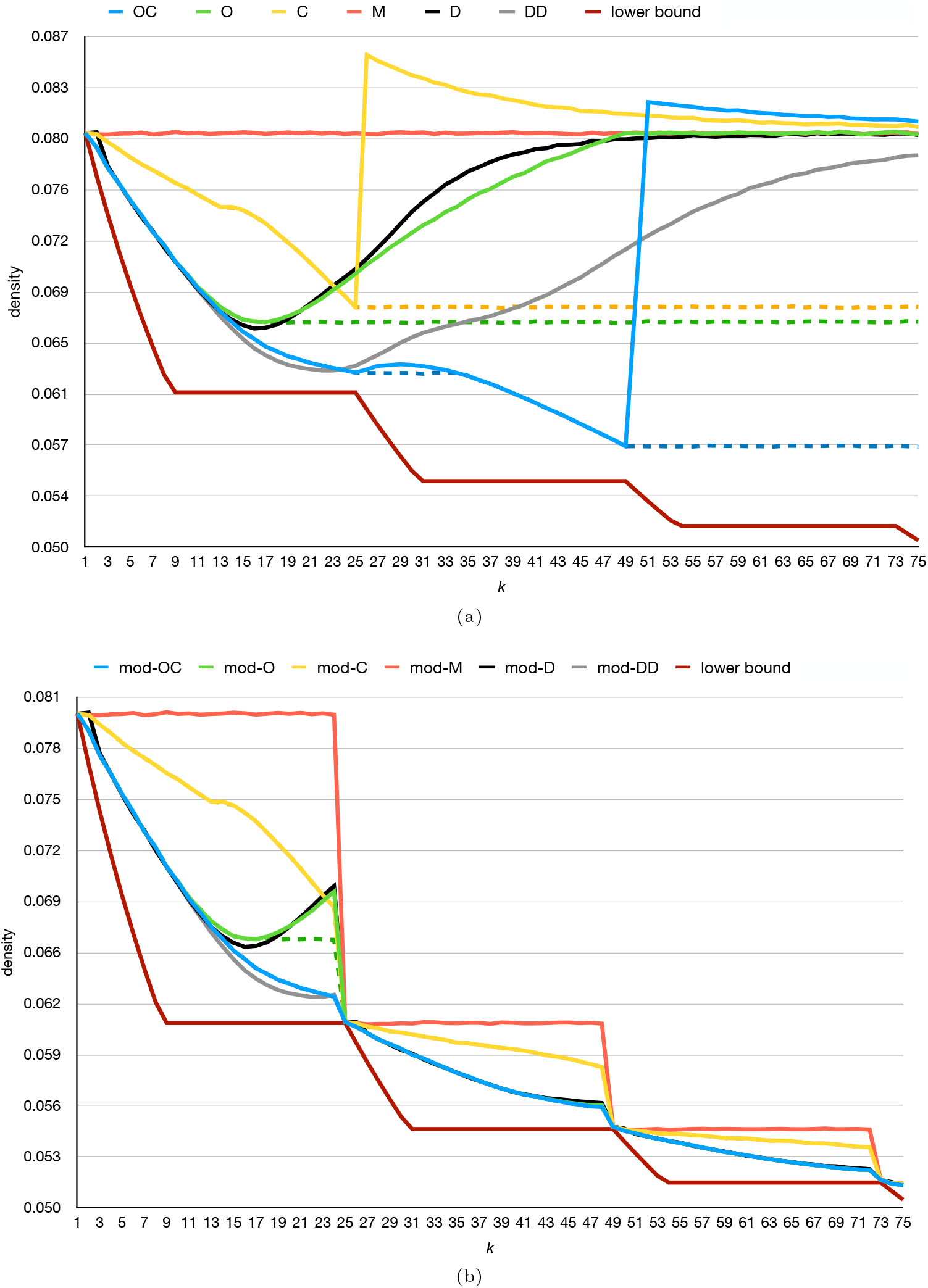
Density for *w* = 24 and by varying *k*, measured on a random string of ten million i.i.d. random characters for *σ* = 256. For all methods that require the parameter *s*, we use *s* = 1 for the solid lines. The dashed lines, instead, use the best choice of *s*. We use *r* = 1 for all methods.

In the legend and remaining text, we use the following abbreviations:

- `M`: the random **m**inimizer.
- `C`: **c**losed syncmer minimizer, corresponding to the miniception;
- `O`: **o**pen syncmer minimizer, where open syncmers are preferred over *k*-mers;
- `OC`: the **o**pen-**c**losed minimizer from Section 3.1;
- `D` and `DD`: the ***d**ecycling* and ***d**ouble **d**ecycling* set based methods introduced by Pellow et al. [11].

For details on the decycling methods, we refer to the original paper and to Section 3 of [12] for a review of the method. These schemes map each *k*-mer to a complex number and prefer those with argument between 0 and 2*π/k*, and in our implementation the arithmetic involved tends to be slightly less efficient than the other discussed methods. Nevertheless, double decycling often has the lowest density of all schemes as also evident from Figure 3.

As apparent, the `OC` scheme performs remarkably better than the other two variants of miniception when *k* approaches *w*, and indeed has a similar shape to the density of the decycling set based methods, `D` and `DD`. In fact, this similarity is even closer when the alphabet is large and *s* = 1, Figure 6a. However, compared to `D` and `DD, OC` is simpler, more intuitive, and even faster to compute as it does not involve arithmetic with complex numbers. All the solid lines in the plot use *s* = 4. The dashed lines are instead obtained by taking the best choice of *s* for each *k*. It is interesting to note that, for very small *k* (say, in the range [5..10]) and the best choice of *s*, the methods `O` and `OC` achieve better density than the decycling set based methods.

#### SSHash indexes

To give a concrete idea of how open-closed minimizers can be useful in practice, we use them to build SSHash indexes [3, 4], across some different datasets. (We remark that the SSHash data structure can use *any* sampling scheme that respects a window guarantee. The default choice in SSHash is to use the classic random minimizer.) We test the chromosome 13 of the human genome, the whole human genome (GRCh38), a small pangenome of 100 *Salmonella Enterica* genomes [30], and the whole genome of the the *Ambystoma Mexicanum* (the “axolotl”), which has one of the largest genomes (more than 18 billion distinct *k*-mers for *k* = 31). Table 2 reports space usage of SSHash in bits/*k*-mer on these datasets, for (*w, k*) = (11, 21): the open-closed minimizer makes SSHash consistently smaller, improving its space usage by at least 11% and up to 14.5%.

##### Algorithm 4

Pseudocode for the *mod-sampling* and the *extended mod-minimizer* methods. For the extended mod-minimizer, *A* : ∑^*w*+*k−*1^ → [*w* + *k* − *t*] is *any* sampling scheme used to define the anchor. We assume that *A* defines an order between *t*-mers and that its definition might use additional parameters to define the sampling, like *s* ≤ *t* in case *A* is the open-closed minimizer from Section 3.1.

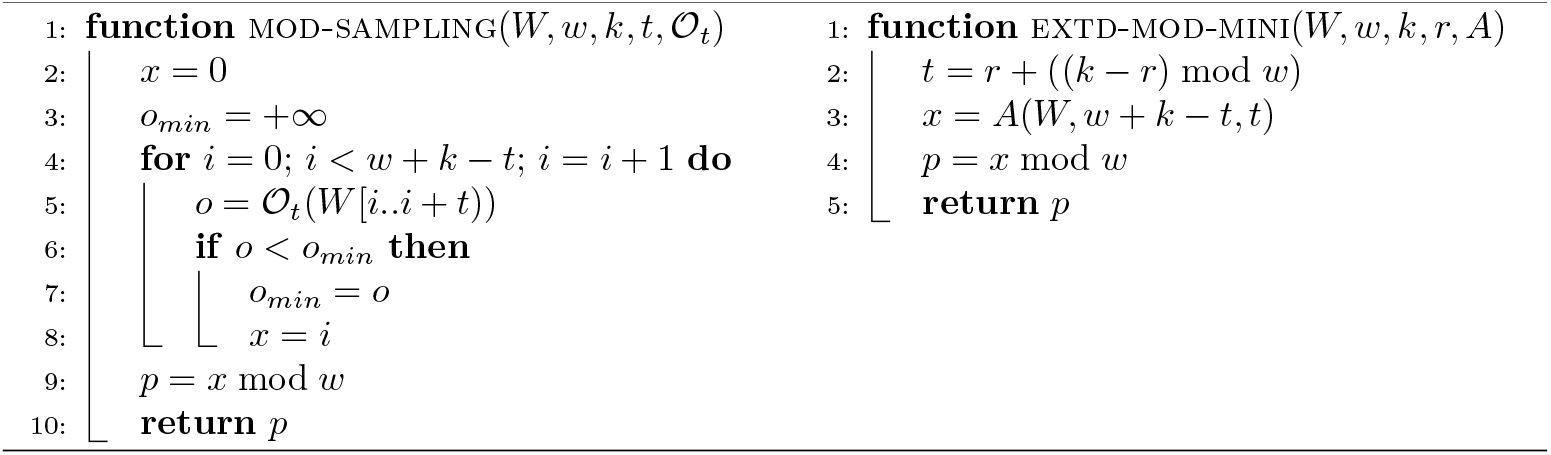

#### Behaviour for large *k*

Lastly in this section, we observe that all methods discussed so far cease to work well when *k* grows (with *s* fixed) and their density worsens towards that of a random minimizer. The reason is that as *k* grows, fewer and fewer *k*-mers are an open/closed syncmer. Thus, more and more windows of *w k*-mers will not contain a single “special” *k*-mer, and thus fall back to the random minimizer. A larger value of *s* can be used to prevent this, but will still not allow density to improve beyond a constant as *k* → ∞.

For example, the C method (miniception^2^) has a sharp increase in the density when *k* − *s > w*, which is exactly when a window is not guaranteed to contain a closed syncmer anymore. A similar effect is observed for the `OC` method, where for *k* − *s >* 2*w* no open nor closed syncmer might be found in a window. For the `O` method, instead, the probability that a window contains no open syncmer already starts to increase already before *k* reaches *w*.

The decycling set based methods (`D, DD`), instead, do not perform well for large *k* as the universal hitting sets contain roughly 1*/k* of *k*-mers and hence become too sparse to ensure most windows contain a *k*-mer in these sets.

To compensate for this drop in effectiveness, we need another method which is the subject of the next section.

## 4 The large-*k* case: the extended mod-minimizer

In this section we consider methods tailored for the case where *k* is larger than *w*, which we refer to as the “large-*k*” case.

Recently, Groot Koerkamp and Pibiri [12] introduced the *mod-sampling* method – a framework to obtain minimizer schemes that have low density when *k > w*. The method is illustrated in Algorithm 4 (left). It simply determines the position *x* of the smallest *t*-mer in the window for some *t* ≤ *k*. It then samples the *k*-mer at position *x* mod *w*. The complexity of the method is clearly *O*(*w* + *k* − 1).

As we argued in Section 1, the method works intuitively well because the smallest *t*-mer acts as an “anchor” for potentially many more than *w* consecutive, making the mod-sampling exhibit a locally optimal behavior when the smallest *t*-mer does not change: either it samples the same *k*-mer from consecutive windows or it samples the *k*-mer that it *w* positions apart from the last sampled *k*-mer. This effect is depicted in Figure 1a.

In this section we extend this method to work with any anchoring mechanism, and not just the smallest *t*-mer found by a random minimizer.

### 4.1 The extended mod-minimizer

We first fix the choice of the parameter *t* for mod-sampling as *t* = *r* +((*k r*) mod *w*), for some lower bound *r* ≤ *t*. (We use *r* = 4 in our experiments.) Groot Koerkamp and Pibiri [12] showed that this choice of *t* minimizes the density of mod-sampling and it gives a minimizer scheme named the *mod-minimizer*. Furthermore, when *r >* (3 + *ε*) log_*σ*_ (*w* + *k* − 1) for some *ε >* 0 and the order 𝒪_*t*_ is random, the density of the mod-minimizer tends to the optimal 1*/w* as *k* → ∞. Kille et al. [14] also showed that, for large alphabets, the mod-minimizer has near-optimal density when *k* ≡ 1 (mod *w*), and not just when *k* is large.

Let us call *anchor of length t* the *t*-mer that is selected by the mod-minimizer to determine the position of the sampled *k*-mer. Here we note that the mod-minimizer can be further extended to consider any arbitrary sampling function *A* : ∑^*w*+*k−*1^ → [*w* + *k* − *t*] to determine the anchor, where *A* can be a minimizer, a more general forward scheme, or even a local scheme. This *extended mod-minimizer* algorithm is shown in Algorithm 4 (right). In fact, while the anchor can simply be determined by taking the random minimizer of length *t* as done in [12], this is just one among many possible choices. For example, we showed in Section 3 that closed and open syncmers improve over random minimizers for small *k*. Thus, it makes sense to use the open-closed minimizer of length *t* as anchor in the extended mod-minimizer. We call this new scheme the *open-closed mod-minimizer*. An example is shown in Figure 4.

This scheme converges to optimal density 1*/w* for *k* → ∞ (see Section 4.2) like the mod-minimizer that uses a random minimizer as anchor but, as we will see in Section 4.3, it achieves even lower density for many practical values of *k*.

### 4.2 Analysis

As explained, the extended mod-minimizer generally works with any anchor *A* : ∑^*w*+*k−*1^ → [*w* + *k* − *t*] that samples a *t*-mer from a window. The following theorem shows how the density of the extended mod-minimizer relates to the density of *A* when *A* is a minimizer scheme.

#### Theorem 1.

*Let A* : ∑^*w*+*k−*1^ → [*w* + *k* − *t*] *be a minimizer scheme that selects the smallest t-mer according to some order* 𝒪_*t*_. *Then the density of the mod-minimizer is given by the probability that A samples a t-mer in a position p* ≡ 0 (mod *w*) *in a context of two consecutive windows, whose total length is w*+*k characters and contains w* + *k* − *t* + 1 *t-mers*.

*Proof*. Consider two consecutive windows *W* and *W*^*′*^ of length *w* + *k* − 1 of a uniform random string. Let *x* and *x*^*′*^ be the position of the smallest *t*-mer in *W* and *W*^*′*^ respectively, and let *p* = *x* mod *w* and *p*^*′*^ = *x*^*′*^ mod *w* be the positions of the sampled *k*-mers. Let *y* ∈ {*x, x*^*′*^ + 1} be the absolute position of the smallest *t*-mer in the two windows.

Since *A* is a forward scheme, we can compute its density as the probability that a difference *k*-mer is sampled from *W* and *W*^*′*^. First note that the two consecutive windows contain a total of *w* + *k*− *t* + 1 *t*-mers, and thus, 0 ≤ *y* ≤ *w* + *k* − *t*, where *w* + *k* − *t* is divisible by *w* since *t* ≡ *k* (mod *w*).

When *y* ≡ 0 (mod *w*), this implies 0 *< y < w* + *k* − *t*, and thus, the two windows share their smallest *t*-mer. Thus, *p* = *x* mod *w* = *y* mod *w* and *p*^*′*^ +1 = *x*^*′*^ mod *w* +1 = (*y* − 1) mod *w* + 1. Since *y* ≢ 0 (mod *w*), this gives *p*^*′*^ + 1 = *y* mod *w*, and thus, the two windows sample the same *k*-mer.

When *y* ≡ 0 (mod *w*), there are two cases. When *y* = *x* (and thus *y < w* + *k* − *t*), we have *p* = *x* mod *w* = *y* mod *w* = 0, and since the *k*-mer starting at position 0 is not part of *W*^*′*^, the second window must necessarily sample a new *k*-mer. Otherwise, we must have *y* = (*x*^*′*^ + 1) ≡ 0 (mod *w*), which implies *p*^*′*^ = *x*^*′*^ mod *w* = (*y* − 1) mod *w* = *w* − 1, and since the *k*-mer starting at position *w* − 1 in *W*^*′*^ is not part of *W*, again the second window must necessarily sample a new *k*-mer.

To conclude, the two windows sample distinct *k*-mers if and only if the smallest *t*-mer occurs in a position *y* ≡ 0 (mod *w*).

Using Theorem 1, we can obtain a closed-form formula for the density of the extended mod-minimizer when *A* is a random minimizer, because we know that the position of the smallest *t*-mer is uniformly distributed in [*w* + *k* − *t* + 1].

#### Lemma 1

([12], Lemma 9). *For any ϵ >* 0, *if t >* (3 + *ϵ*) log_*σ*_ (*w* + *k* − 1), *the probability that a random window of w* + *k* − *t t-mers contains two identical t-mers is o*(1*/*(*w* + *k* − 1)), *which tends to 0 for k* → ∞.

#### Theorem 2.

*If t* ≡ *k* (mod *w*) *satisfies the condition in Lemma 1 and A is the random minimizer, then the density of the extended mod-minimizer is*

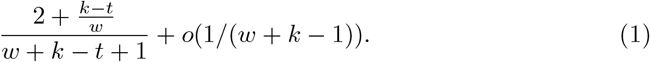

*When w is fixed and k* → ∞, *the density tends t o* 1*/w*.

*Proof*. By Theorem 1, we must bound the probability that a position *y* ≡ 0 (mod *w*) is sampled in a context of *w* + *k* − *t* + 1 *t*-mers. When the smallest *t*-mer in the context is unique, its position is uniformly distributed. Since there are 1 + (*w* + *k* − *t*)*/w* positions *y* such that *y* ≡ 0 (mod *w*), the probability is 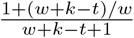. Otherwise, we can bound the probability that a non-unique smallest *t*-mer is in such a position by *o*(1*/*(*w* + *k* − 1)) by Lemma 1. We directly obtain the result. It is immediate to see that the density goes to 1*/w* when *w* is fixed and *k* → ∞.

For different anchors *A*, the position of the smallest *t*-mer may not be uniformly distributed in {0, …, *w* + *k* − *t*}, and hence they may induce a different density for the extended mod-minimizer.

#### Computing the density of the open-closed mod-minimizer

While we do not have a closed-form formula for the probability that *A* samples a *k*-mer at position *p* ≡ 0 (mod *w*) when *A* is an open-closed minimizer, we can extend the analysis made in Section 3.2 for the open-closed minimizer to the open-closed mod-minimizer. By Theorem 1, we must compute the probability that the sampled *k*-mer is in a position *p* ≡ 0 (mod *w*). Thus, we change the definition of *charged k-mer* to not only be the leftmost and rightmost *k*-mer, but to include any *k*-mer at a position *p* ≡ 0 (mod *w*). Apart from accounting for these additional charged *k*-mers, the recursive algorithm shown in Algorithm 3 stays the same.

### 4.3 Density

Figure 5 shows the density of the same methods compared in Figure 3, but when they are used as anchors (of length *t* = *r* +(*k* − *r*) mod *w*, for *r* = 4) for the extended mod-minimizer. Thus, we prefix their names by “mod”. The plot shows that the extended mod-minimizer can be used as a method to “lift” any method from small to large *k*, i.e., to improve its density when *k > w*. Indeed, all methods have better density than the random mod-minimizer (method mod-M) and, among those, the open-closed mod-minimizer (method mod-OC) should be preferred over mod-DD for reasons already discussed. Figure 6b in Appendix B shows the equivalent plot of Figure 5 for *σ* = 256, with similar results.

Considering again Table 2, the open-closed mod-minimizer consistently improves over the open-closed minimizer from Section 3 by 3 − 4%, resulting in a decrease in SSHash’s space of 18% on the whole human genome. We stress that this improvement in space usage is obtained *without* modifying the SSHash data structure, but only by changing the sampling algorithm.

## 5 Conclusions and future work

In this work, we introduced the open-closed minimizer, a method that achieves very low density for the case when *k* ≤ *w*. Technically speaking, this is achieved by extending the miniception method of Zheng et al. [10] to also sample open syncmers (and not just closed syncmers). This is based on the intuition that open syncmers should be preferred over closed syncmers as they satisfy a distance lower bound, i.e., that two consecutive open syncmers tend to be well apart from each other. This new method thus achieves density that is practically as low as the double decycling method by Pellow et al. [11], but is simple, intuitive, and computationally efficient.

Then, we also extended the mod-minimizer by Groot Koerkamp and Pibiri [12], that works by selecting the smallest *t*-mer inside the window and uses this “anchor” to determine the position of the *k*-mer to sample. We extended this method to consider *any* arbitrary sampling scheme to select the *t*-mer. This can yield better densities than the original mod-minimizer, depending on the choice of the anchor. The extended mod-minimizer can thus be used to improve the density of *any* method when *k > w*. For example, by combining the open-closed minimizer with the mod-minimizer we obtained the so-called *open-closed mod-minimizer*, which achieves even lower density in practice than the random mod-minimizer for *k* where it is not already provably optimal.

To show the direct impact of these results, we replaced the random minimizer used in the SSHash data structure [3, 4] with the open-closed mod-minimizer. This simple change decreases the space usage by up to 18%, e.g., on the whole human genome. As future work, it would also be interesting to quantify the impact of our new schemes on other applications such as read mapping. For example, our sampling schemes could be used in the minimap2 software [31].

### Future work

The analysis of the extended mod-minimizer is more complicated than the version using random minimizers and, currently, we lack closed-form formulas for its density. Future work could therefore try to derive such formulas (or tight approximations) for specific anchors, like the open-closed minimizer.

More generally, one could investigate how much closer to the 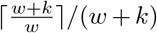 lower bound by Kille et al. [14] schemes can get. In particular, forward schemes having density equal to the lower bound must never sample overlapping *k*-mers. Compared to random minimizers, where *k*-mers can overlap by *k* − 1, open syncmers get halfway there by being roughly *k/*2 positions apart. Preliminary results suggest that when *k* is small (up to ≈ *w/*6) it is indeed possible to design a scheme where sampled *k*-mers usually do not overlap, and hence to achieve density very close to the lower bound. Thus, the schemes introduced in this paper should not be considered as a definitive word on this important matter.

## Author contributions

R.G.K.: Conceptualization, Methodology, Software, Validation, Writing – Review and Editing; D.L.: Conceptualization, Software; G.E.P.: Conceptualization, Methodology, Software, Validation, Writing – Original Draft. All authors read and approved the final manuscript.

## Declarations

None to report.

## A Probability distributions

Table 3 and Table 4 report the probability distribution for the count configurations (*C, C*_*c*_) and (*O, C, O*_*c*_, *C*_*c*_) respectively, as used in the analysis from Section 3.

## B Density plots for σ = 256

Figure 6a and Figure 6b show the same density plots as, respectively, Figure 3 and Figure 5, but for a the larger alphabet *σ* = 256. In this case, we therefore used *s* = 1. Note how all methods exactly match the lower bound by Kille et al. [14] for *k* ≡ 1 (mod *w*).

## Footnotes

1 It was recently shown that the density is slightly below 2*/*(*w* +1) when *k ≥ w* and *w* is not too small [24].

2 Zheng et al. [10] proved that the density achieved by miniception can be upper bounded by 1.67*/w* + *o*(1*/w*), when *k* ≥ *w* and *s* = *k − w* + 1. But even then, the density is only dependent on *w* and therefore cannot improve when *k* grows.

## References

[1] Ndiaye, M., Prieto-Baños, S., Fitzgerald, L.M., Yazdizadeh Kharrazi, A., Oreshkov, S., Dessimoz, C., Sedlazeck, F.J., Glover, N., Majidian, S.: When less is more: sketching with minimizers in genomics. Genome Biology 25(1), 270 (2024) 10.1186/s13059-024-03414-4

[2] Zheng, H., Marçais, G., Kingsford, C.: Creating and using minimizer sketches in computational genomics. Journal of Computational Biology 30(12), 1251–1276 (2023) 10.1089/cmb.2023.0094

[3] Pibiri, G.E.: Sparse and skew hashing of k-mers. Bioinformatics 38, 185–194 (2022) 10.1093/bioinformatics/btac245

[4] Pibiri, G.E.: On weighted k-mer dictionaries. Algorithms for Molecular Biology 18(1), 1–20 (2023) 10.1186/s13015-023-00226-2

[5] Marchet, C., Kerbiriou, M., Limasset, A.: BLight: Efficient exact associative structure for k-mers. Bioinformatics 37(18), 2858–2865 (2021) 10.1093/bioinformatics/btab217

[6] Fan, J., Khan, J., Singh, N.P., Pibiri, G.E., Patro, R.: Fulgor: A fast and compact k-mer index for large-scale matching and color queries. Algorithms for Molecular Biology 19(1), 1–21 (2024) 10.1186/s13015-024-00251-9

[7] Pibiri, G.E., Fan, J., Patro, R.: Meta-colored compacted de Bruijn graphs. In: Research in Computational Molecular Biology - 28th Annual International Conference, RECOMB 2024, Cambridge, MA, USA, April 29 - May 2, 2024, Proceedings, pp. 131–146 (2024). 10.1007/978-1-0716-3989-49

[8] Roberts, M., Hayes, W.B., Hunt, B.R., Mount, S.M., Yorke, J.A.: Reducing storage requirements for biological sequence comparison. Bioinformatics 20(18), 3363–3369 (2004) 10.1093/bioinformatics/bth408

[9] Schleimer, S., Wilkerson, D.S., Aiken, A.: Winnowing: local algorithms for document fingerprinting. In: Halevy, A.Y., Ives, Z.G., Doan, A. (eds.) Proceedings of the 2003 ACM SIGMOD International Conference on Management of Data, San Diego, California, USA, June 9–12, 2003, pp. 76–85 (2003). 10.1145/872757.872770

[10] Zheng, H., Kingsford, C., Marçais, G.: Improved design and analysis of practical minimizers. Bioinformatics 36, 119–127 (2020) 10.1093/bioinformatics/btaa472

[11] Pellow, D., Pu, L., Ekim, B., Kotlar, L., Berger, B., Shamir, R., Orenstein, Y.: Efficient minimizer orders for large values of k using minimum decycling sets. Genome Research 33(7), 1154–1161 (2023) 10.1101/gr.277644.123

[12] Groot Koerkamp, R., Pibiri, G.E.: The mod-minimizer: A Simple and Efficient Sampling Algorithm for Long k-mers. In: 24th International Workshop on Algorithms in Bioinformatics (WABI 2024), vol. 312, pp. 11–11123 (2024). 10.4230/LIPIcs.WABI.2024.11

[13] Marçais, G., DeBlasio, D.F., Kingsford, C.: Asymptotically optimal minimizers schemes. Bioinformatics 34(13), 13–22 (2018) 10.1093/bioinformatics/bty258

[14] Kille, B., Koerkamp, R.G., McAdams, D., Liu, A., Treangen, T.J.: A near-tight lower bound on the density of forward sampling schemes. bioRxiv (2024) 10.1101/2024.09.06.611668

[15] Edgar, R.: Syncmers are more sensitive than minimizers for selecting conserved k-mers in biological sequences. PeerJ 9 (2021) 10.7717/peerj.10805

[16] Shaw, J., Yu, Y.W.: Theory of local k-mer selection with applications to long-read alignment. Bioinformatics 38(20), 4659–4669 (2022) 10.1093/bioinformatics/btab790

[17] Pike, G., Alakuijala, J.: CityHash. https://github.com/aappleby/smhasher/blob/master/src/City.cpp (2011)

[18] Belazzougui, D., Botelho, F.C., Dietzfelbinger, M.: Hash, displace, and compress. In: Algorithms - ESA 2009, 17th Annual European Symposium, Copenhagen, Denmark, September 7–9, 2009. Proceedings. Lecture Notes in Computer Science, vol. 5757, pp. 682–693 (2009). 10.1007/978-3-642-04128-061

[19] Pibiri, G.E., Trani, R.: Parallel and external-memory construction of minimal perfect hash functions with pthash. IEEE Transactions on Knowledge and Data Engineering 36(3), 1249–1259 (2024) 10.1109/TKDE.2023.3303341

[20] Mohamadi, H., Chu, J., Vandervalk, B.P., Birol, I.: ntHash: recursive nucleotide hashing. Bioinformatics 32(22), 3492–3494 (2016) 10.1093/bioinformatics/btw397

[21] Chikhi, R., Limasset, A., Medvedev, P.: Compacting de Bruijn graphs from sequencing data quickly and in low memory. Bioinformatics 32(12), 201–208 (2016) 10.1093/bioinformatics/btw279

[22] Khan, J., Kokot, M., Deorowicz, S., Patro, R.: Scalable, ultra-fast, and low-memory construction of compacted de bruijn graphs with cuttlefish 2. Genome biology 23(1), 190 (2022) 10.1186/s13059-022-02743-6

[23] Cracco, A., Tomescu, A.I.: Extremely fast construction and querying of compacted and colored de Bruijn graphs with GGCAT. Genome Research 33, 1198–1207 (2023) 10.1101/gr.277615.122

[24] Golan, S., Shur, A.M.: Expected density of random minimizers. arXiv (2024) 10.48550/arXiv.2410.16968

[25] Orenstein, Y., Pellow, D., Marçais, G., Shamir, R., Kingsford, C.: Designing small universal k-mer hitting sets for improved analysis of high-throughput sequencing. PLoS computational biology 13(10), 1005777 (2017) 10.1371/journal.pcbi.1005777

[26] Ekim, B., Berger, B., Orenstein, Y.: A randomized parallel algorithm for efficiently finding near-optimal universal hitting sets. In: Schwartz, R. (ed.) Research in Computational Molecular Biology - 24th Annual International Conference, RECOMB 2020, Padua, Italy, May 10-13, 2020, Proceedings. Lecture Notes in Computer Science, vol. 12074, pp. 37–53 (2020). 10.1007/978-3-030-45257-53

[27] Kille, B., Garrison, E., Treangen, T.J., Phillippy, A.M.: Minmers are a generalization of minimizers that enable unbiased local jaccard estimation. Bioinformatics 39(9), 512 (2023) 10.1093/bioinformatics/btad512

[28] Loukides, G., Pissis, S.P., Sweering, M.: Bidirectional string anchors for improved text indexing and top-k similarity search. IEEE Transactions on Knowledge and Data Engineering 35(11), 11093–11111 (2023) 10.1109/TKDE.2022.3231780

[29] Marçais, G., Pellow, D., Bork, D., Orenstein, Y., Shamir, R., Kingsford, C.: Improving the performance of minimizers and winnowing schemes. Bioinformatics 33(14), 110–117 (2017) 10.1093/bioinformatics/btx235

[30] Rossi, M., Silva, M.S.D., Ribeiro-Gonçalves, B.F., Silva, D.N., Machado, M.P., Oleastro, M., Borges, V., Isidro, J., Viera, L., Halkilahti, J., Jaakkonen, A., Palma, F., Salmenlinna, S., Hakkinen, M., Garaizar, J., Bikandi, J., Hilbert, F., Carriço, J.A.: INNUENDO whole genome and core genome MLST schemas and datasets for Salmonella enterica (2018) 10.5281/zenodo.1322563

[31] Li, H.: Minimap2: pairwise alignment for nucleotide sequences. Bioinformatics 34(18), 3094–3100 (2018) 10.1093/bioinformatics/bty191

